# Metatranscriptomic Insights into Host-Microbiome Interactions Underlying Asymptomatic COVID-19 Cases

**DOI:** 10.1101/2025.08.18.670804

**Authors:** Sanjana Fatema Chowdhury, Md. Murshed Hasan Sarkar, Syed Muktadir Al Sium, Showti Raheel Naser, Md. Saddam Hossain, Md. Ahashan Habib, Shahina Akter, Tanjina Akhtar Banu, Barna Goswami, Iffat Jahan, Tanay Chakrovarty, Md. Maruf Ahmed Molla, Tasnim Nafisa, Mahmuda Yeasmin, Asish Kumar Ghosh, Md. Salim Khan

## Abstract

In recent years, the research on Coronavirus disease 2019 (COVID-19) has surged rapidly due to its infectious nature and epidemiological importance as a pandemic. Association of other pathogenic microbes or different human microbiomes with COVID-19 severity is well established. While much is known about COVID-19, studies exploring how host-pathogen interactions at the transcriptomic level influence disease severity are still limited. This research focuses on metatranscriptomic perspective of COVID-19 patients from different Bangladeshi cohorts. Metatranscriptomic sequencing was performed using the extracted RNA of forty different nasopharyngeal samples. After preprocessing and assembly of the genome sequence data, taxonomic identification and diversity along with antibiotic resistance pattern was analyzed using different bioinformatic pipelines. COVID-19 positive and asymptomatic positive patients had a higher abundance of pathogenic and multidrug resistant bacteria whereas the Healthy or recovered patients had higher fungal population. Differential gene expression analysis was also performed to identify the upregulated and downregulated genes responsible for different biological and immunological pathways of humans. Here, immunological response related genes were mostly upregulated in positive cases which was also evident in the proinflammatory cytokines upregulation. Moreover, asymptomatic positive cases showed low TLR-4 expression, which is key to recognizing pathogen-associated molecular patterns (PAMPs) and triggering pro-inflammatory immune responses. This study can clarify the gene expression and signaling pattern of COVID-19 patients with different severity. This may also suggest that some populations exhibit reduced basal expression of TLR-4, which may suppress innate immune activation following infection and contribute to asymptomatic clinical outcomes; however, this hypothesis requires further investigation. Future studies should aim to validate these associations using larger, ethnically diverse cohorts with comprehensive clinical and demographic data.

## Introduction

Severe acute respiratory syndrome coronavirus 2 (SARS-CoV-2), the causative agent of coronavirus disease 2019 (COVID-19), has created a catastrophic event throughout the world known as COVID-19 pandemic [1,2]. The novel virus first emerged from Wuhan, China in late December 2019 as an indicator of numerous cases of serious pneumonia and has been considered as one of the deadliest human pathogens after Spanish Flu [3,4].

However, fatality and prognosis of COVID-19 is influenced by several factors such as, patient’s age or preexisting comorbidities (diabetes, hypertension, asthma etc.) [5,6]. Vaccines have made it possible to reduce the severity of the disease, but emergence of such viral disease still remains a threat. Since the COVID-19 outbreak, there has been a significant increase in the collection of epidemiological, clinical, and immunological data. But the molecular mechanism underlying SARS-CoV-2 virus pathogenesis and prognosis was still not explored much [7,8]. This is because a thorough understanding of the pathogenesis of COVID-19 should enable the development of effective strategies to enhance the generation of effective anti-SARS-CoV-2 immune responses.

The respiratory microbiota consists of a wide range of bacteria and is distributed in the upper respiratory tract (URT). This microbial flora contains either commensals or pathobionts. Various investigations have shown that the severity of SARS-CoV-2 infection is likely to be influenced by the co-existing bacteria, fungi, and archaea in the respiratory tracts [9,10]. The SARS-CoV-2 virus enters the body by inhalation and binds to epithelial cells in the nasal cavity. It then replicates and migrates through the respiratory tract and gradually proceeds towards the lungs to initiate infection which in turn mounts a vigorous innate immune response for clearing up the virus [11]. So, it is believed that during this propagation and immune response, the microbiota residing in the respiratory airways may be altered, and inclusion of some of the pathobionts might alleviate the course and severity of the disease caused by SARS-CoV-2 virus [11].

Nowadays, Next-generation sequencing (NGS) approaches, such as metagenomics, is the widely used method for analyzing what kind of microbial population is present in a sample. But metagenomics mainly focuses on the genomic content and identification of microbes present within a community, while metatranscriptomics, a subset of metagenomics, provides the diversity of the active genes within such community, their expression level profile and how these levels change with alterations in environmental conditions. Thus, to understand the role of respiratory microbiome in COVID-19 severity and to elucidate host gene expression with functional profiling metatranscriptomics approaches are preferable.

In this study, nasopharyngeal samples of different severity groups of COVID-19 patients were collected and subjected to sequencing to analyze both respiratory microbiome and differential gene expression and immunological signaling pathways of the host [12,13]. This information not only can be used for taxonomic characterization of known or novel microbes and host response analysis but also determines the functional profile of biological processes of normal and altered respiratory microbiome [13,14].

## Methodology

### Sampling and Transcriptome Sequencing

Nasopharyngeal samples from forty different people were collected for the study. COVID-19 positive and suspected individuals of different categories were included in the study. The cohort had individuals from ICU, COVID-19 positive, negative, reinfected, recovered, and asymptomatic positive patients as well. Asymptomatic positive patients were healthcare staff who had to go through routine checkup of COVID-19 RT-PCR test. However, the samples were broadly divided into four different classes for the sake of analysis, and they are: Severe, Mild, Asymptomatic, and Negative (S1 file) The samples were collected using universal transfer media (UTM) and RNA was extracted using the ReliaPrep™ Viral TNA Miniprep System, Promega (USA) as soon as the samples arrived at the laboratory. Illumina NextSeq550 was used for sequencing purposes, and the sequencing library was prepared using TruSeq RNA Library Preparation Kit as per the manufacturers protocol. Preprocessing and alignment were performed using RNA Seq Alignment app software.

### Microbiome diversity and antibiotic resistance analysis

Different microbial population (Prokaryotes and Eukaryotes) present in the nasopharyngeal samples were checked using different R packages (version 4.2), to determine the association of COVID-19 severity with microbial diversity. “Phyloseq” [15] and “Vegan” [16] were used to conduct normalization procedures via rarefaction and to assess disparities in abundances. These packages facilitated the comprehensive analysis of diversity and composition across various sample groups. Alpha diversity (differences within the sample groups) metrics were assessed employing the Wilcoxon Rank Signed test, while PERMANOVA, utilizing the Adonis test with 999 permutations (implemented through the Set.seed function), was employed to evaluate discrepancies in beta diversity. Furthermore, for the graphical representation of diversity measures and compositional analysis, R packages such as “ggpubr” and “ggplot2” were employed [17].

Antibiotic resistant genes of the data were checked using CZID [18] platform. A Circos plot was generated using “circlize” R package [19] based on the number of hits on each resistant gene.

### Microbial metatranscriptomics, host differential gene expression and enrichment analysis

Microbial metatranscriptomics analysis was performed using Microbiome Metatranscriptomics pipeline (version 10.2) from BaseSpace, Illumina. The raw data was plotted in heatmap using Pheatmap package in R. Differential gene expression of the host genome is important to determine the functional changes of human body due to diverse microbial community. DESeq2 package [20] was used for differential gene expression analysis and several other bioinformatic tools were used for gene enrichment analysis and protein-protein interaction prediction.

### miRNA expression analysis

Differential expressions of miRNA was analyzed using BioJupies [21] among different sample groups. Then the upregulated and downregulated miRNAs were plotted in Venn diagram separately to identify the common and unique miRNA among different groups. R package ggvenn was used for this purpose. The case specific miRNAs were then used for analyzing different gene enrichment analysis such as Reactome pathway, KEGG signaling pathway, metabolic pathway, gene ontology etc. using RNAenrich webserver [22].

### Signaling pathway analysis

A software package Pathview [23] was used for pathway-based data integration and visualization. Pathview generates native KEGG pathway graphs, hence is natural and more readable for humans. Here different colors indicate the most upregulated and downregulated genes. Important immunological pathways were analyzed and compared among different study groups for finding associations using different statistical analysis.

## Results

### Taxonomic and microbial diversity analysis

Alpha diversity measured using Shannon and Simpson Diversity metrics showed significant species diversity within the sample groups in terms of prokaryotes (Fig 1). But in terms of Eukaryotes, the box and whisker plot show widespread distribution of species diversity for the severe group, whereas for the other three groups, the data were clustered.

**Fig 1:**
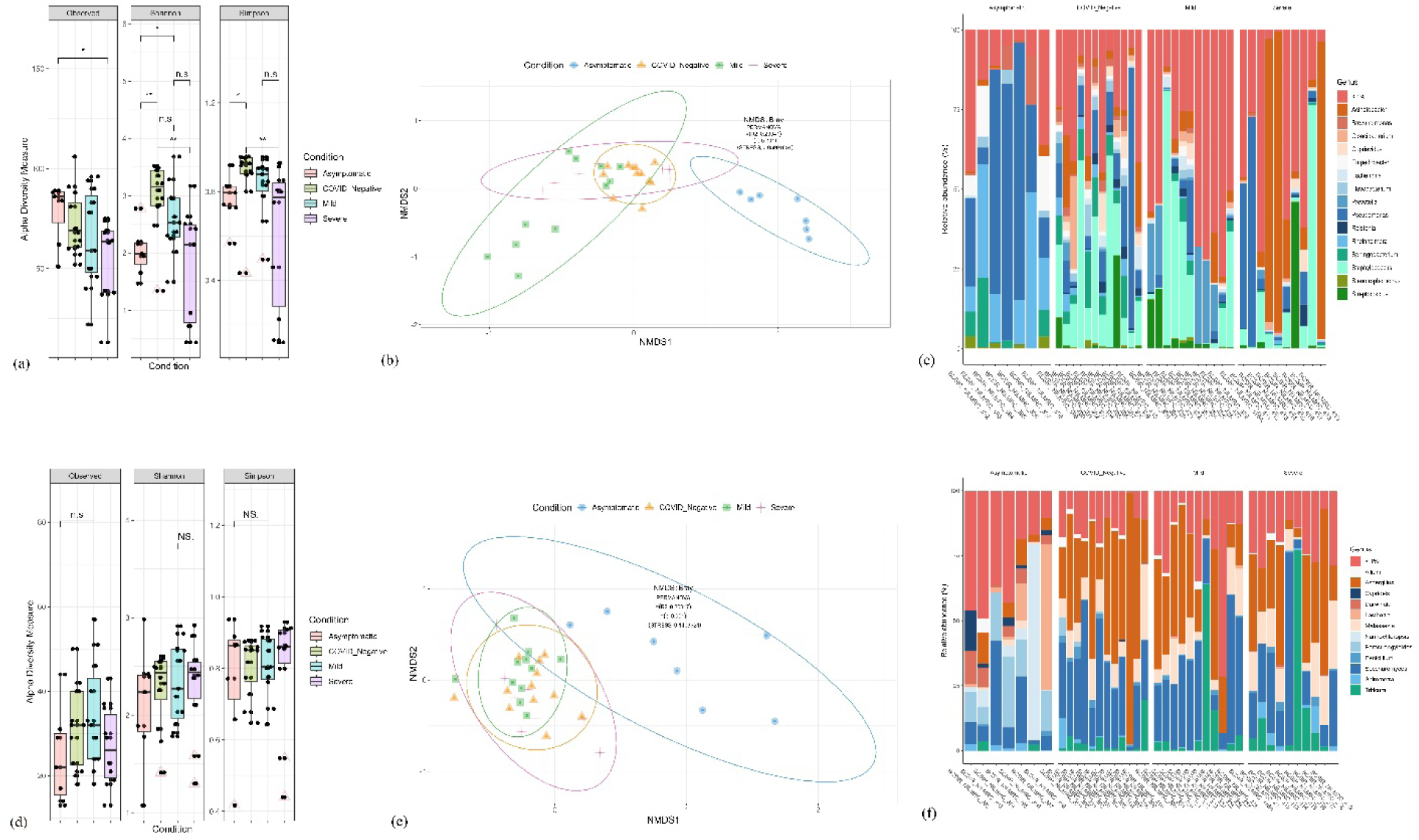
Alpha (a) and Beta (b) diversity indexes in Prokaryotes. Alpha (d) and Beta (e) diversity indexes in Eukaryotes. Genus level Relative abundance of four major study cohorts. Here, (c) Prokaryotes and (f) Eukaryotes.

However, Bray–Curtis distance method Beta diversity analysis showed a separate cluster for Asymptomatic group in PCoA but the clusters of other three sample groups (Negative, Mild, Severe) overlapped for both the prokaryotes and eukaryotes (Fig 1).

The relative abundance of taxonomic analysis among the four major study groups showed the high abundance of pathogenic and multidrug resistant bacterial strain among COVID-19 positive patients irrespective of severity (Fig 1c,1f). Moreover, COVID-19 positive patients had fungal dominance apart from SARS-CoV-2 viral infection.

### Microbial metatranscriptomics and antimicrobial resistance analysis

The diversity in microbial community delved into looking more closely at the condition of microbial metatranscriptomics in the samples. This analysis reveals the functions of microbial communities by checking the transcriptome of the microbiome in each environment.

Differences in gene expression patterns and functional activity were observed in different study cohorts of COVID-19 patients in terms of severity. Substantial up-regulation of translation machinery and ribosomal proteins, including multiple 50S and 30S ribosomal proteins, was observed in the microbial gene expression profile of asymptomatic patients. Along with that, genes linked to porins, and cold shock proteins were also highly expressed. Moreover, Gene Ontology (GO) analysis revealed enrichment in molecular functions including DNA binding, GTPase activity, and magnesium ion binding, as well as biological processes like translation elongation, protein transport, and the tricarboxylic acid cycle.

Significant up-regulation of viral genes, such as replicase polyprotein 1AB, spike glycoprotein, and RNA-directed RNA polymerase, was observed in mild COVID-19 patients. The structural components of virions, host immune response regulation, and processes related to the viral life cycle were found to have enrichment, according to GO analysis.

Stress response proteins, such as Rubredoxin and class D beta-lactamase, were up-regulated in the microbial gene expression profile of severe COVID-19 patients. GO analysis showed inhibition of the host antiviral signal, modification of host protein ubiquitination, and enrichment in processes associated with the viral life cycle.

Genes related to beta-lactamase activity, cytochrome C oxidase activity, and aerobic respiration were all overexpressed in the healthy group. GO enrichment reflected normal cellular activity in the absence of severe viral infection and was linked to host immunological suppression by viral processes, sensitivity to drugs, and mitochondrial function.

Analysis of antimicrobial resistance gene (ARG) profiles across the study cohorts revealed a markedly higher abundance and diversity of ARGs in the asymptomatic COVID-19 group compared to mild, severe, and negative cohorts. The asymptomatic COVID-19 cohort harbored a distinct resistome, enriched in genes conferring resistance to aminoglycosides, trimethoprim, phenicols, and rifamycins, which were largely absent in other severity groups. However, the β-lactamase gene *TEM1-D* was present across all cohorts and highlighted in red in the circos plot to denote its high abundance (Fig 2c).

**Fig 2:**
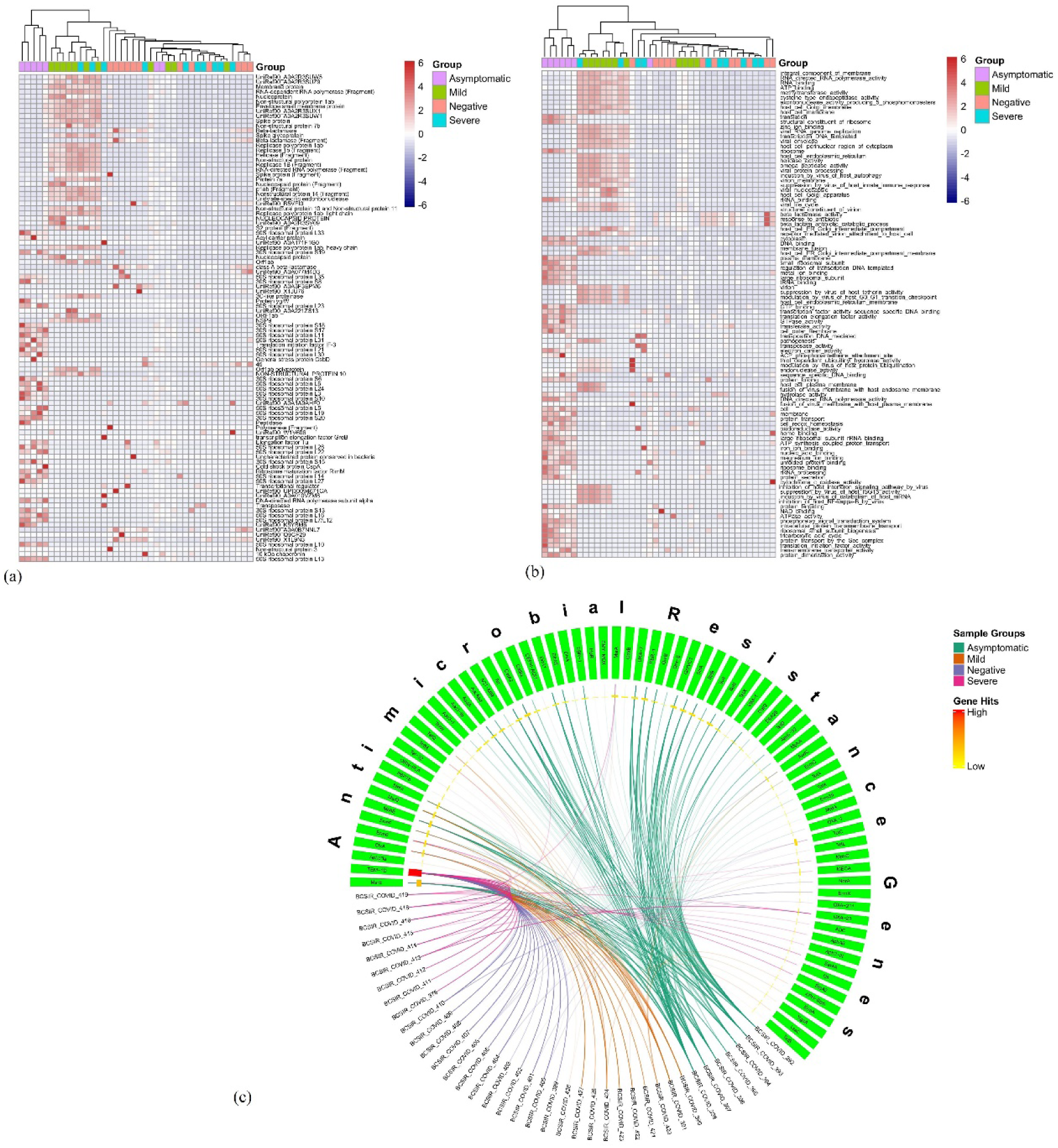
Here, (a) showing Gene Family abundance and (b) showing GO Term abundance in all the samples. Also, (c) showing a circos plot analysis of antibiotic resistance in asymptomatic positive patients visualizing the presence of different AMR genes across the isolates. A heatmap in the inner circle of the plot illustrates the frequency of each AMR gene across all isolates, with red indicating high frequency and yellow indicating low frequency.

### COVD-19 associated host gene expression and gene enrichment

The different transcriptional patterns of the Severe, Mild, Healthy, and Asymptomatic groups may be clearly seen in the PCA plot (Fig 3). The biological significance of the transcriptome variations is shown by the clustering patterns and confidence ellipses, which show the variability within groups and the gap between them. Host gene expression analysis results were further plotted on different groups for better understanding of the outcomes.

**Fig 3:**
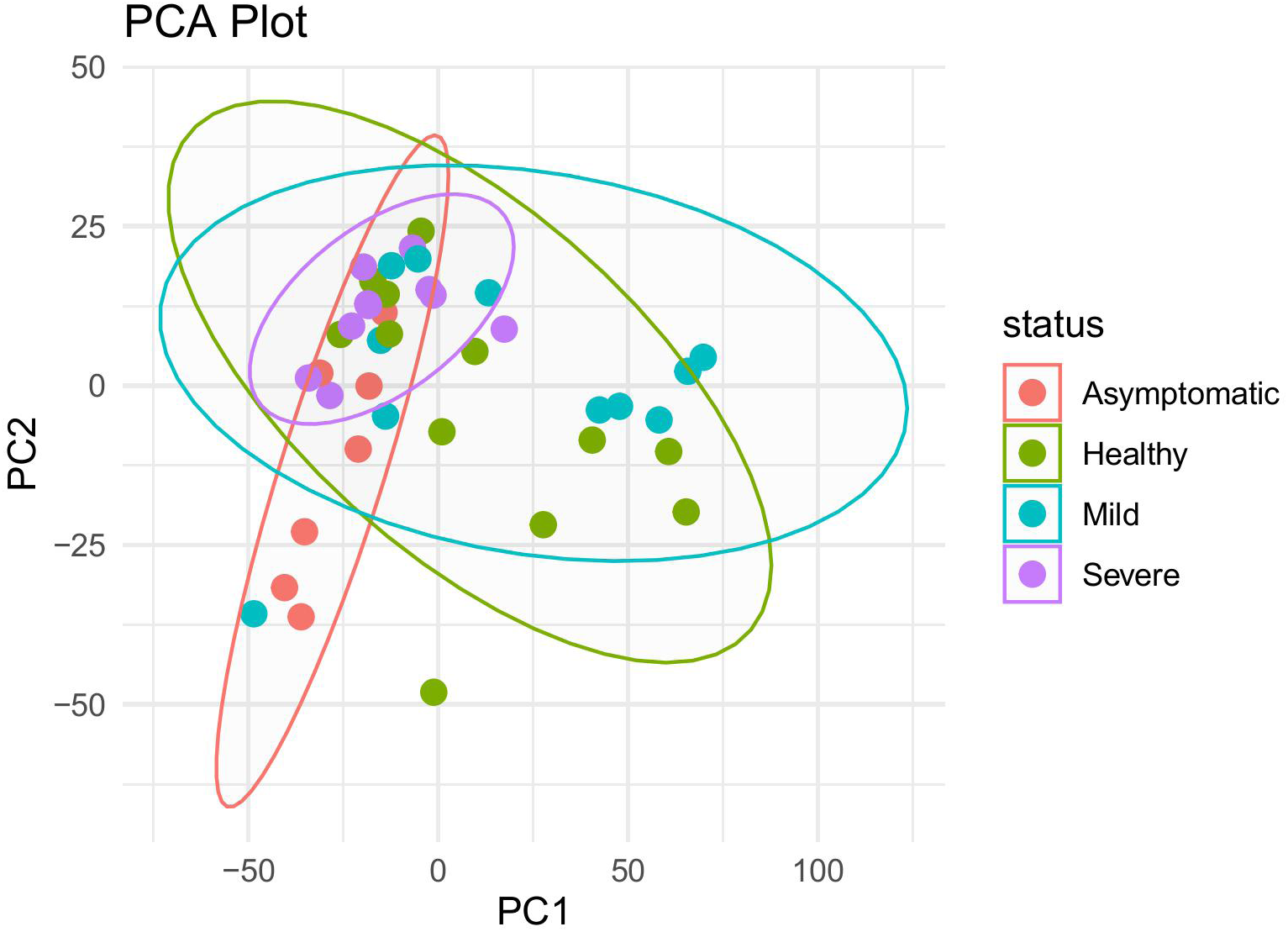
PCA plot showing the overlapping clusters of host gene expression data

Differential expressions of mild and negative patients showed 23 upregulated genes in the mild group than negative patients. The top five expressed genes, such as, ENSG00000229976 (lncRNA), ENSG00000087274 (ADD1), ENSG00000185745 (IFIT1), ENSG00000119922 (IFIT2), ENSG00000169245 (CXCL10) are marked in Fig 4. After the enrichment analysis of the upregulated genes, it was evident that mostly virus associated immunological genes were upregulated in COVID-19 positive patients. Cellular response to cytokine stimulus, innate immune response, response to virus, cytokine mediated signaling pathway, Type-1 interferon signaling pathway etc. were most prominent ones. This analysis followed a protein-protein interaction network analysis to ensure the association among the pathways. Immune response related genes formed a cluster among themselves according to that analysis and binding regulation gene can be seen distant than the cluster.

**Fig 4:**
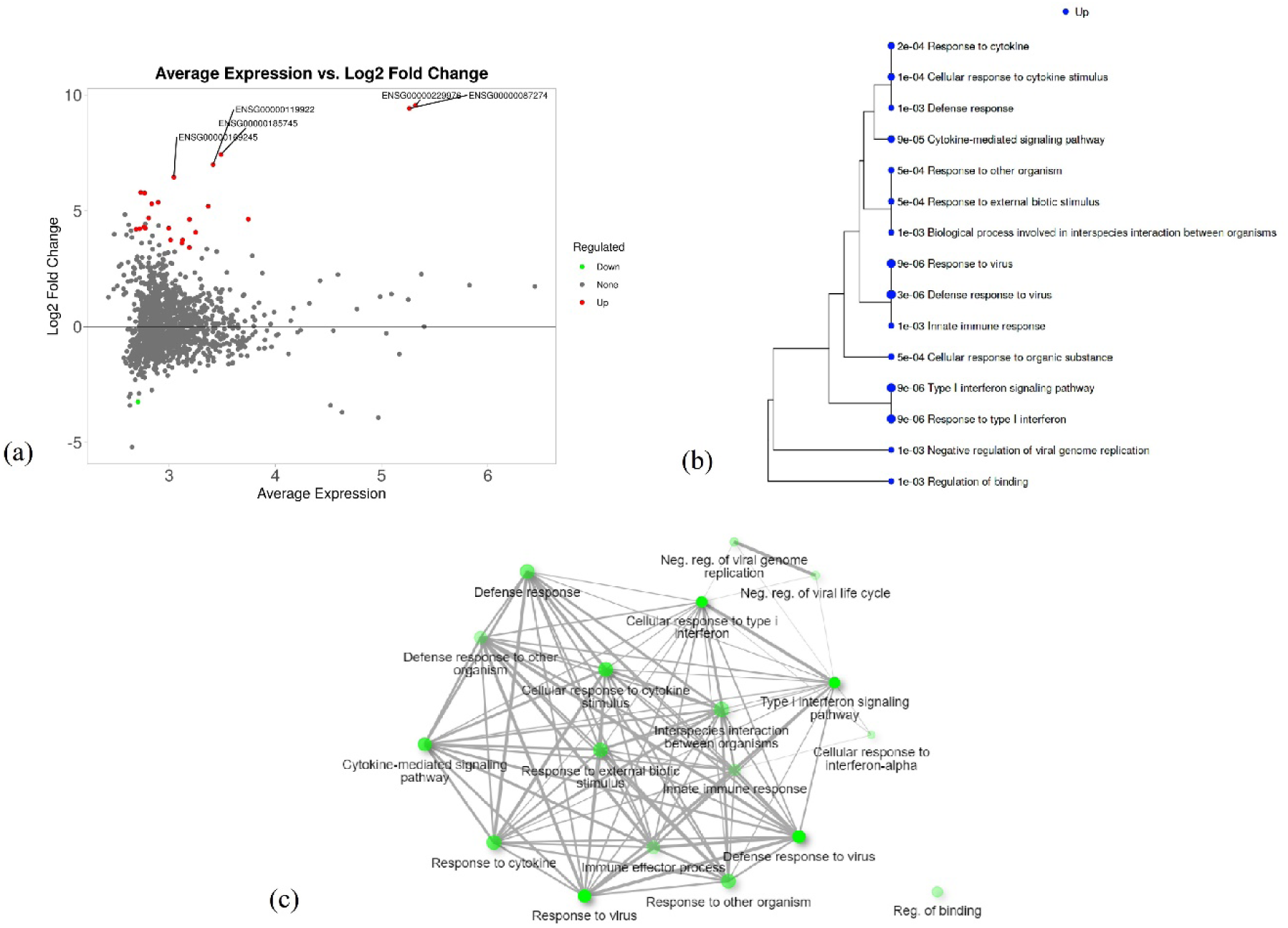
Differential gene expression analysis between Mild and Negative groups. Fig 4(a) shows MA plot for the differential gene expression of this group. Fig 4(b) showing enrichment analysis of the upregulated genes and Fig 4(c) showing PPI network among them.

Moreover, differential expression analysis of mild vs asymptomatic positive group and symptomatic (Mild and Sever) vs asymptomatic positive group was performed. The mild and asymptomatic positive group showed 23 downregulated genes and symptomatic vs asymptomatic group showed 19 downregulated genes. The top expressed genes from both groups are shown in the MA plot. ENSG00000198522 (GPN1), ENSG00000160293 (VAV2), ENSG00000135111 (TBX3), ENSG00000151458 (ANKRD50), ENSG00000230987. Further enrichment analysis of the downregulated genes showed 16 common genes, when plotted in Venn diagram. These genes showed physiological functional processes in gene ontology enrichment, and they all fall into one cluster upon network analysis among themselves.

Furthermore, COVID-19 negative group showed 12 downregulated genes than the asymptomatic group. The upregulated genes are ENSG00000210082 (MT-RNR2), ENSG00000211459 (MT-RNR1) and the top expressed downregulated genes are as follows, ENSG00000130032 (PRRG3), ENSG00000198522 (GPN1), ENSG00000087274 (ADD1), ENSG00000229976 (lncRNA), ENSG00000287389 (lncRNA). The GO biological process enrichment analysis showed downregulation of immunological signaling genes in negative patients.

### COVID-19 influences the immunological signaling pathway

KEGG pathway analysis of different cohorts showed significant results depending on severity. Further analysis on these KEGG pathways, especially immunological signaling pathways, were done to check on more detailed effects.

Pathview analysis on several immunological KEGG pathways showed significant differences among COVID-19 positive, negative, asymptomatic positive and recovered patients.

COVID-19 positive patients, irrespective of the severity, showed upregulated inflammatory response with the release of pro-inflammatory cytokines like IL-1β, IL-12 and resulted in a cytokine storm of IL-8, IL-12, MCP-1, IP-10. However, when COVID-19 negative samples were compared with asymptomatic positive samples, inflammatory cytokines were comparatively downregulated. But the cytokines were upregulated when the order was reversed i.e., asymptomatic vs negative.

An interesting pattern was found with IL-6 which is the most important cytokine for any viral infection. In COVID-19 positive cases, IL-6 was found to be moderately downregulated, which is unusual. But when the cytokine-cytokine receptor signaling was checked, IL-6 receptor was highly upregulated. Furthermore, MMP-1 was also upregulated in positive cases, which is reported to be increased with the upregulation of IL-6. As nasopharyngeal swab was used as the sample, cytokine level changes were not evident that much. If blood samples were taken, the results would be more meaningful.

Chemokines are an integral part of cytokine storm after a viral infection. Reports suggest association of CCL2, CCL3, CCL4, CCL7 and CXC subfamily with the severity of cytokine storm. These chemokines were upregulated in our COVID-19 positive cases as well. Asymptomatic positive patients showed a slightly different expression pattern of chemokines, especially the receptors and CXC subfamily, in comparison with COVID-19 negative patients.

TLRs play a major role in the initiation of innate immune responses, with the production of inflammatory cytokines, type-1 IFN and other mediators. TLR4 stimulation induced the strongest effect in terms of cytokine release.

Here in this study, asymptomatic positive patients showed downregulation of TLR4 in comparison with negative samples. As TLR4 plays a major role in the initiation of cytokine storm, it may be suggested from this data that, downregulation of TLR4 may be responsible for their asymptomatic behavior.

The miRNA based Reactome pathway analysis showed downregulation of IL-4 and IL-13 signaling in Asympotomatic vs Negative individuals. The mild vs healthy group and severe vs healthy group didn’t show any significant miRNA expression profile.

## Discussion

The interconnection between human microbiome and disease development has already been studied well by the scientific community [24–26]. Microbiome composition is shaped by various environmental factors, and in turn, influences host health and immune function. During the COVID-19 pandemic, one of the major questions arose regarding the underlying factors contributing to variable disease severity. While most of the research proved the association of comorbidity, age, sex, and immunosuppression [27–29], the potential influence of the human microbiome has also garnered attention. In particular, the microbiome’s role in explaining the high prevalence of asymptomatic infections, often critical in silent transmission chains, remains an area of active investigation.

In this study, nasopharyngeal samples of different severity groups of COVID-19 patients were subjected to metatranscriptomic analysis for a better understanding of taxonomic and functional profile. Unbiased metatranscriptomic data helps to effectively characterize the microbiome and host response in COVID-19.

Alpha diversity measured using Shannon and Simpson Diversity metrics showed significant species diversity within the sample groups in terms of prokaryotes. But widespread distribution was observed in terms of Eukaryotes (Fig 1d), suggesting greater heterogeneity among these individuals, potentially reflecting increased opportunistic fungal colonization, which has been commonly reported in severe COVID-19 cases [11]. However, between diversity analysis of different groups created a separate cluster for Asymptomatic group for both the prokaryotes and eukaryotes, highlighting the unique species diversity in the group. The rest of the groups showed a closer clustering, suggesting a more stable eukaryotic community.

The tendency to acquire more pathogenic bacteria increases with severity. From the Relative Abundance (RA) analysis, it’s evident that pathogenic strains like *Acinetobacter* and *Pseudomonas* were significantly high in severe cases. Evidence showed the fungal association with the COVID-19 disease, which can sometimes affect the mortality or immune response in the host body [30,31]. In some cases, people also get fungal infection after recovering from viral diseases [32].

According to González. R et al., microbiomes have a significant and broad influence in shaping their host biology, specifically in the immunity and host defense responses [33]. It directly impacts the outcomes of infections by pathogens. This is also reflected in the microbiome metatranscriptomic analysis of different severity in this study. The asymptomatic group of patients did not show any immune response related gene expression rather than showing regular cellular function related genes. This implies a minimum amount of viral interference and an active microbial response. Moreover, the severe and mild group showed a substantial rise of immune response. The data revealed significant mechanisms pertaining to the inhibition of host type I interferon signaling and the stimulation of host autophagy, which are indicative of both viral reproduction in motion and a more regulated immune response. Additionally, there was evidence of elevated cellular stress responses and widespread viral subversion of host cellular pathways and molecular processes. Moreover, the detection of β-lactamase activity in the GO term enrichment analysis indicates not only the presence but also the functional expression of ARGs. This is further supported by the circos plot, where high-intensity signals, particularly from TEM1-D, confirm its prevalence across all severity groups, while unique ARG signatures were enriched in the asymptomatic cohort. Such resistome variation may influence host-microbiome interactions, potentially modulating immune responses and contributing to reduced symptom manifestation [34,35].

However, different research works have shown how the respiratory microbiome is changing with the COVID-19 disease. In this study, COVID-19 related differential host gene expression revealed distinct transcriptional signatures across the study cohorts. COVID-19 positive patients had twenty-three upregulated genes than the negative patients. The upregulated genes were related to responses to viral infection as cytokine storm is already reported to be associated with SARS-CoV-2 infection [36,37]. Further analysis demonstrated the highly interconnected protein-protein interaction network among the upregulated genes.

Additionally, sixteen frequently downregulated genes were identified based on the differentially expressed genes shown in the Venn diagram (Fig 5c) in order to determine the genetic signatures unique to asymptomatic positive patients. These genes were mainly linked to physiological processes, especially those related to reproduction and development, which might be a reflection of underlying pre-existing health conditions in these people.

**Fig 5:**
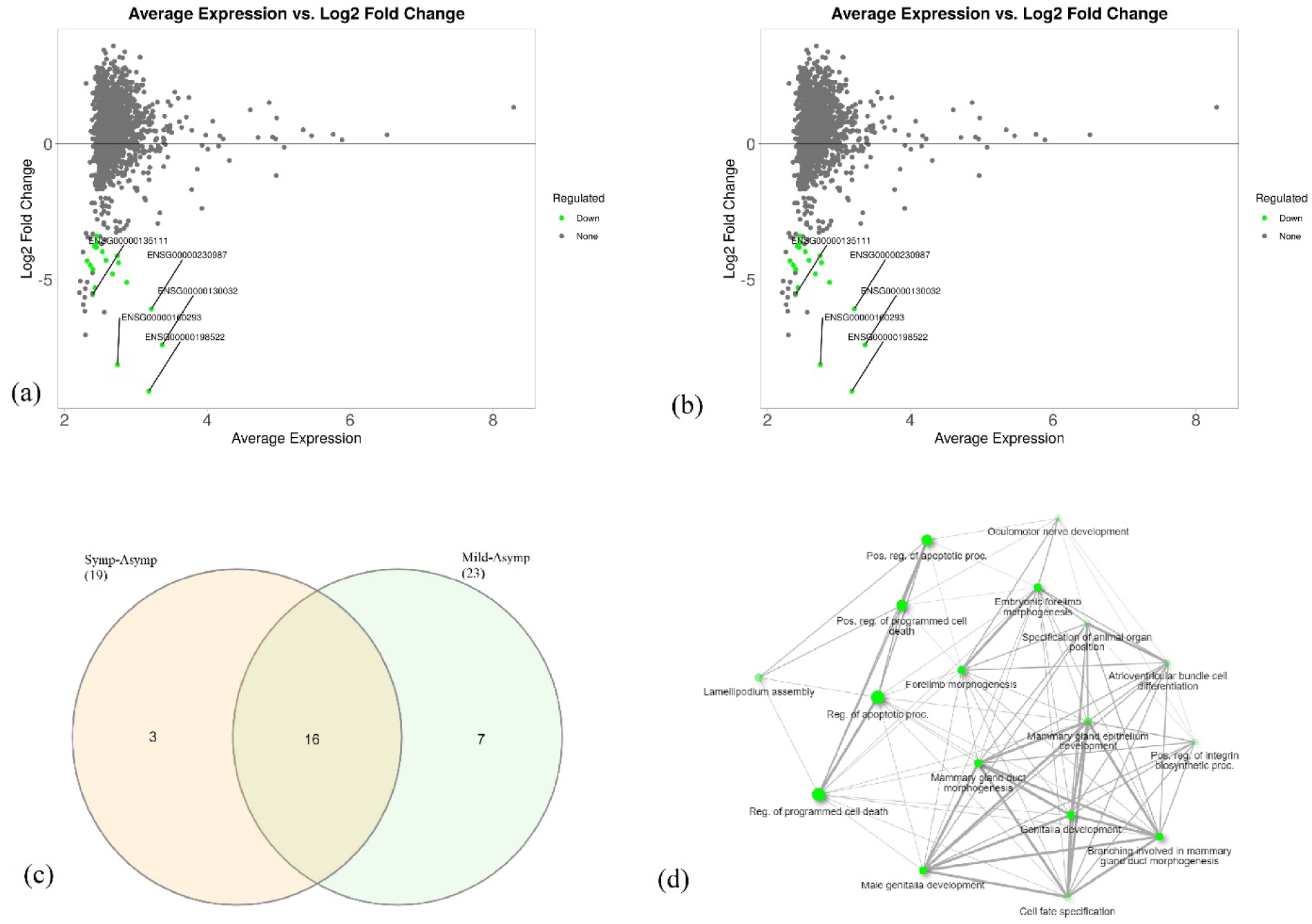
MA plot showing differential gene expression in two different groups. Fig 5(a) shows mild vs asymptomatic group. The top five expressed genes such as, ENSG00000198522 (GPN1), ENSG00000160293 (VAV2), ENSG00000135111 (TBX3), ENSG00000151458 (ANKRD50), ENSG00000230987 are marked here. Fig 5(b) shows symptomatic vs asymptomatic group. Top five expressed genes such as, ENSG00000198522 (GPN1), ENSG00000160293 (VAV2), ENSG00000130032 (PRRG3), ENSG00000230987 and ENSG00000135111 (TBX3) are marked here. Fig 5(c) and Fig 5(d) show the network among the common biological processes.

In addition, differential expression analysis of COVID-19 negative and asymptomatic positive group showed downregulation of immunological signaling genes in negative patients (Fig 6). As they were not affected with the SARS-CoV-2 virus infection, their viral responses were comparatively lower than the other.

**Fig 6:**
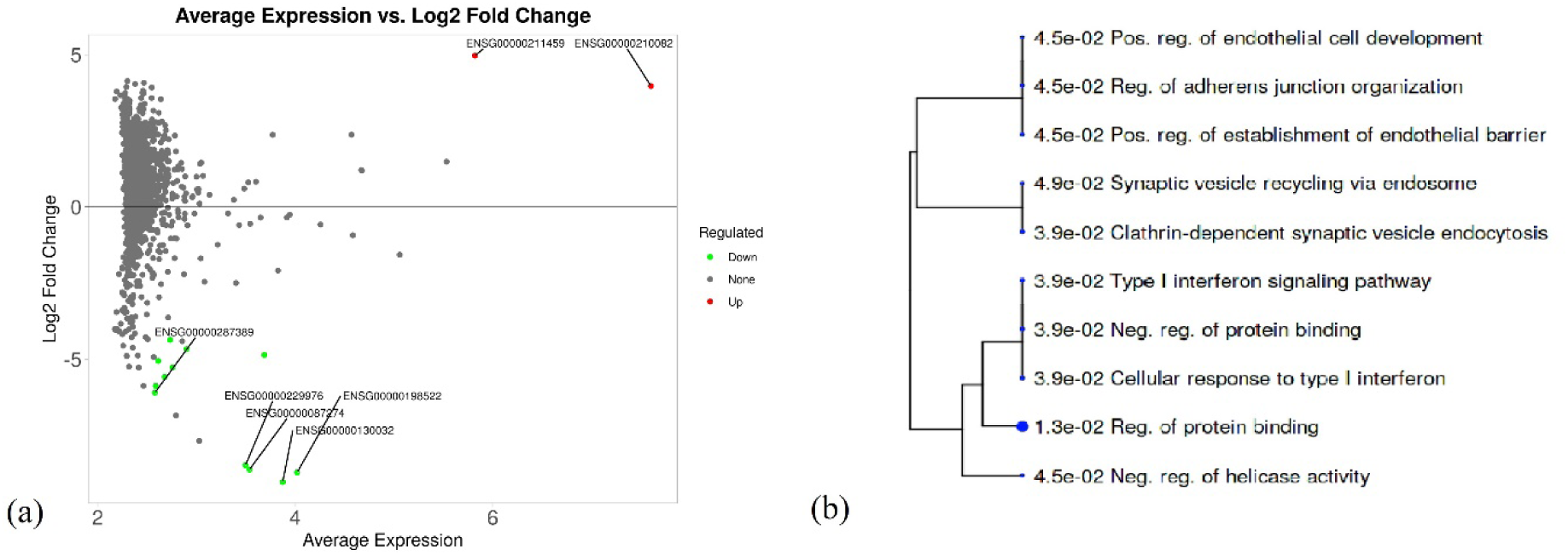
(a) MA plot showing differential gene expression between COVID-19 negative and asymptomatic patients. (b) showing the downregulated genes

**Fig 7:**
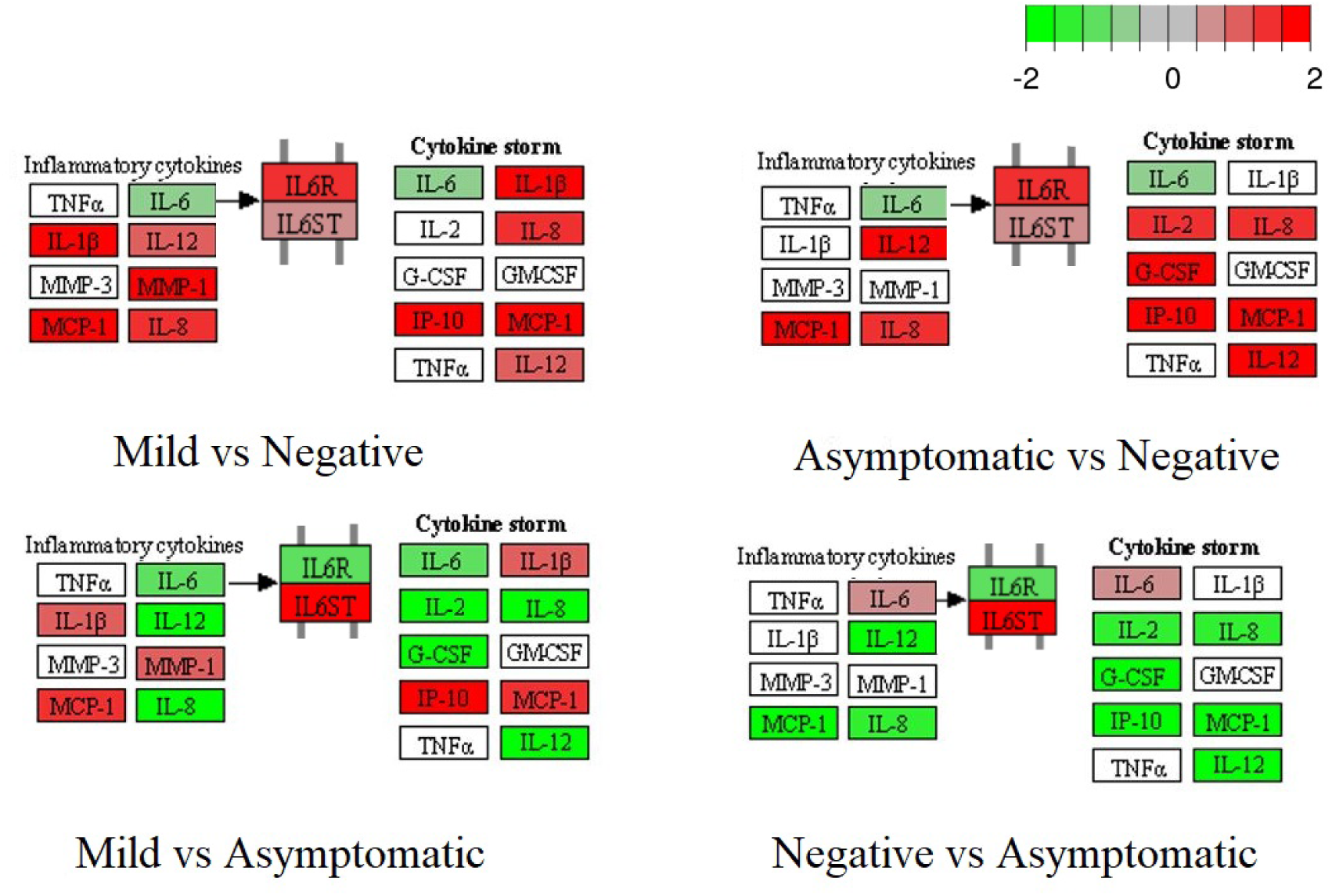
Changes in cytokine signaling in different study cohorts. Here bright red indicates the most upregulated, bright green, the most downregulated.

**Fig 8:**
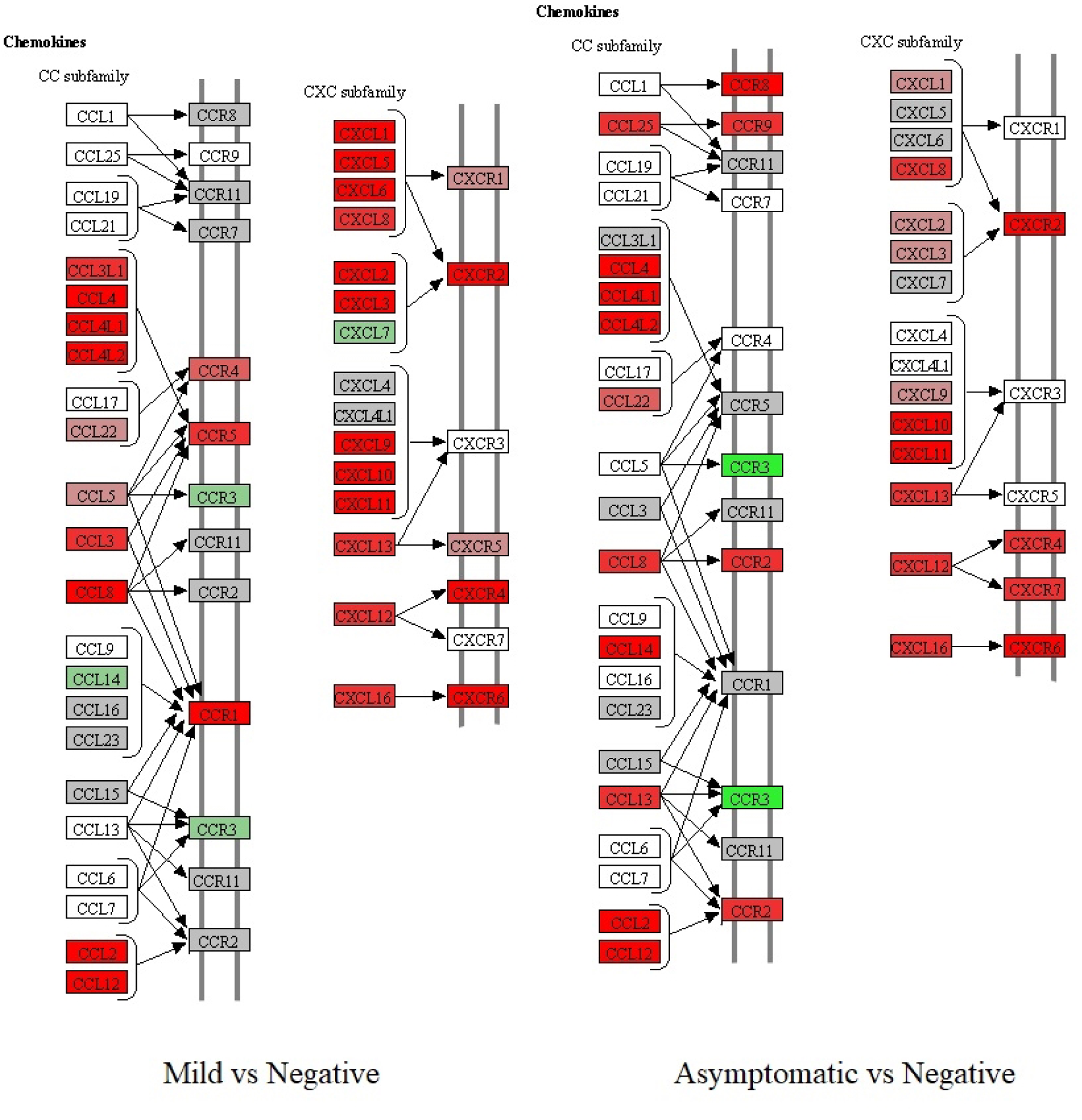
Changes in chemokines and their receptor signaling in different study cohorts. Here bright red indicates most upregulated, bright green, most downregulated.

**Fig 9:**
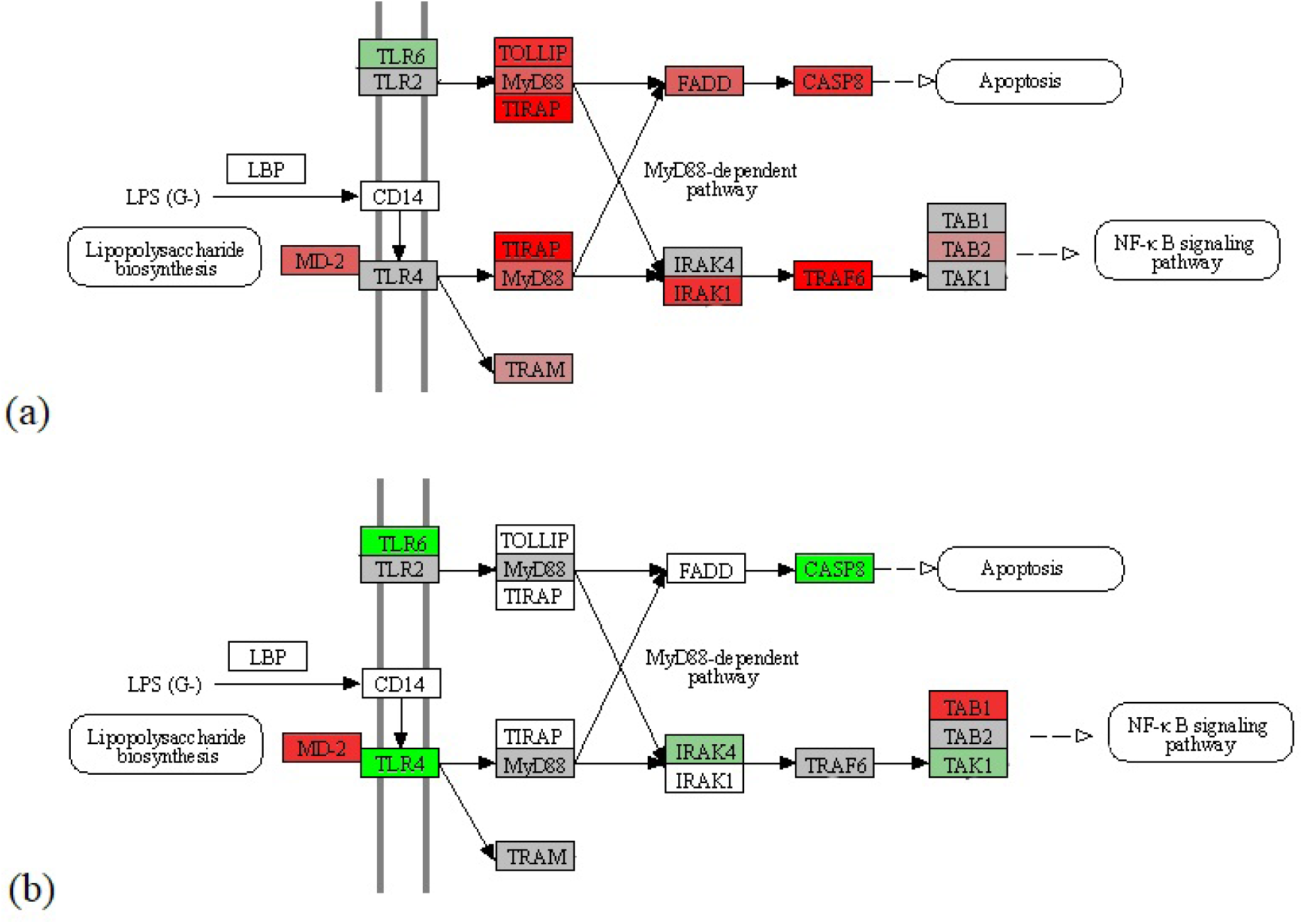
Partial demonstration of Toll like receptor signaling pathway in different groups. (a) Mild vs Negative and, (b) Asymptomatic Positive vs Negative

**Fig 10:**
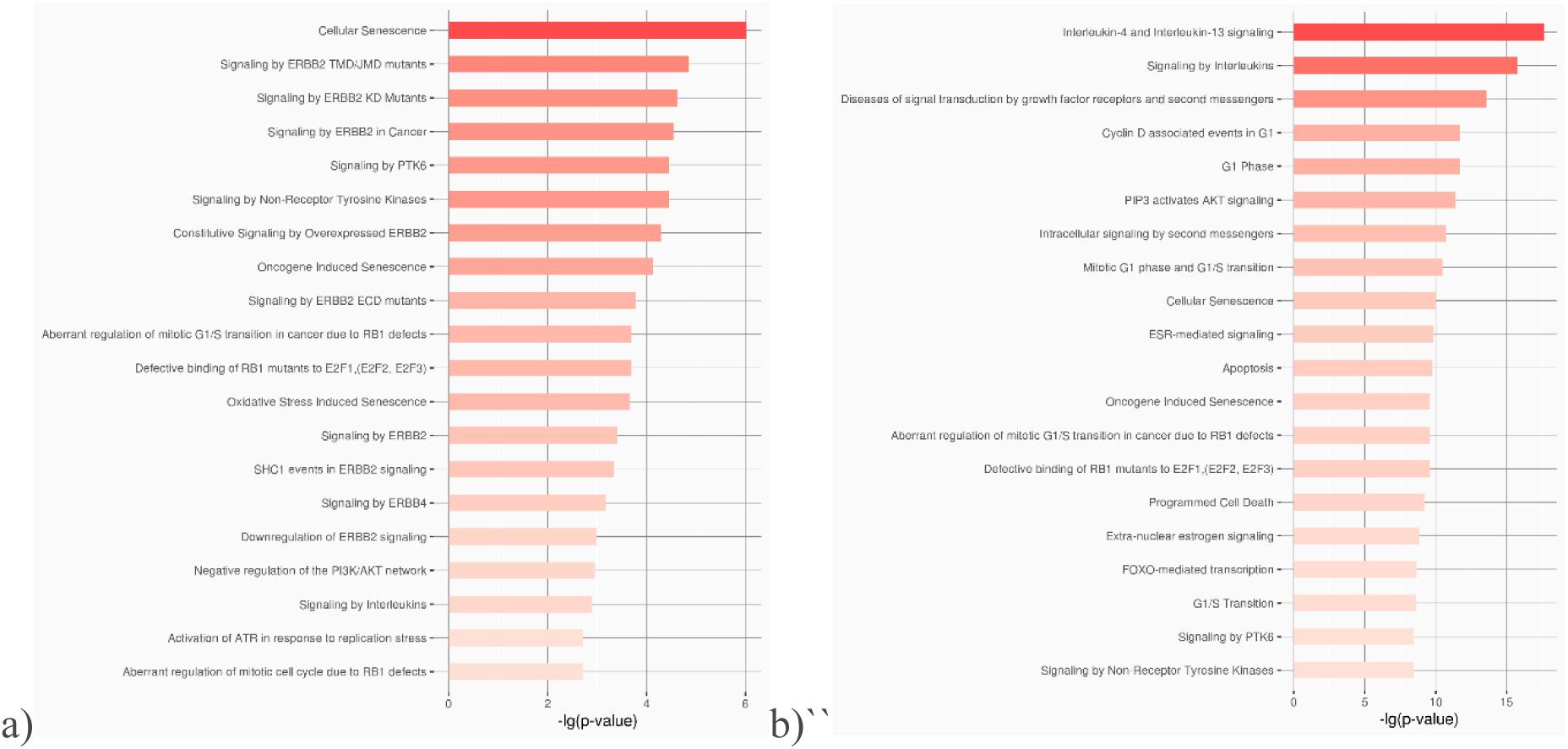
miRNA enrichment analysis on reactome pathway database. Here a) shows the upregulating miRNA in Asymptomatic vs Negative group and b) shows the downregulating miRNA in Asymptomatic vs Negative group.

KEGG pathway analysis is widely used for understanding the biological relevance of the disease or infection with the host body. Different studies have demonstrated the molecular landscape and biomarkers associated with SARS-CoV-2 viral infections using this method [38,39] but in this study a number of immunological signaling was focused on understanding the host response to the infection. Upon entry into the host and subsequent dissemination into the bloodstream, SARS-CoV-2 triggers the activation of immune cells, leading to the release of various pro-inflammatory cytokines. This, in turn, facilitates the release of inflammatory cytokines; in some cases, an exaggerated immune response leads to a cytokine storm, contributing to disease severity and multi-organ dysfunction [40]. In agreement with the available reports on “cytokine storm”, this study also found out the upregulation of some pro-inflammatory cytokines IL-1β and IL-12, which in turn upregulated cytokine storm of IL-8, IL-12, MCP-1, IP-10. In addition, asymptomatic positive samples showed upregulated cytokines against COVID-19 negative samples. The reason here lies in the threshold level of the expression [41]. As the negative samples are not affected by SARS-CoV-2, they are not expected to exhibit a cytokine storm. Though the asymptomatic positive cases may not show symptoms, they are affected by the virus. Thus, they have slight cytokine upregulation. However, this upregulation remains below the threshold needed to trigger symptoms, possibly due to internal regulatory mechanisms. Same goes for the cytokines in asymptomatic positive cases. Chemokines are reported to be involved in COVID-19 progression and severity [42] which was also seen in this study. CCL2, CCL3, CCL4, CCL7 and CXC subfamily were upregulated in the COVID-19 positive cases.

Toll-like receptors (TLRs) are vital components of the innate immune system, functioning as pattern recognition receptors that detect conserved microbial structures and initiate host defense mechanisms [43]. To date, 13 mammalian TLRs (TLR1–TLR13) have been identified, each exhibiting distinct ligand specificities and pattern recognition capabilities [44]. Though TLR4 expression is majorly dependent on lipopolysaccharide (LPS), some studies reported that it is correlated with viral disease severity [45]. This expression mediates the infection symptoms as well. Here, TLR4 has lower expression in asymptomatic positive cases than healthy individuals which answers the questions of these patients no symptom development.

Moreover, the miRNA enrichment analysis of Asymptomatic vs Healthy group showed downregulation of two crucial interleukins, IL-4 and IL-13 which is known to be related with TH-2 mediated immune response [46]. These interleukins are also responsible for inflammation response. Thus, the downregulation of these two major interleukins also supports asymptomatic behavior.

In conclusion, this study reveals distinct microbiome and host transcriptomic profiles across COVID-19 severity groups, with asymptomatic cases showing unique microbial diversity, reduced immune activation, and lower TLR4 expression. Downregulation of pro-inflammatory cytokines and interleukins in this group suggests intrinsic immune regulation may limit symptom development. The detection of active antimicrobial resistance genes further underscores the need for monitoring secondary pathogens in COVID-19. These findings highlight the value of integrating microbiome and host transcriptome analyses to better understand variable disease outcomes, warranting validation in larger, diverse cohorts.

## Conclusion

Viral pathogenesis, in general, encompasses a wide range of complex processes which influence disease severity. Understanding these signaling processes is essential to get an insight into human responses to disease. Asymptomatic COVID-19 cases were a concern for many reasons during the pandemic period, not only due to their potential for silent transmission but also because of the possible long-term effects of the virus on the host. The immunological perspective of asymptomatic COVID-19 cases, found from this study will shed light in understanding other infectious diseases in coming days. Nonetheless, integrating metadata more systematically with genomic and immunological wet-lab data would provide a more comprehensive and accurate picture of these interactions.

## Declaration

### Ethical approval and consent to participate

The ethical permission of the protocol for sample collection from patients, sample processing, and other consecutive laboratory work was taken from the National Institute of Laboratory Medicine and Referral Center (NILMRC) of Bangladesh.

### Availability of data and material

All the data related to this manuscript is available in the supplement files and all the raw data is uploaded to NCBI under the accession number of PRJNA1298058.

### Authors’ contributions

**Experimental Design:** Murshed Hasan Sarkar, Sanjana Fatema Chowdhury, Salim Khan

**Sample Collection:** Md. Maruf Ahmed Molla, Tasnim Nafisa, Mahmuda Yeasmin, Asish Kumar Ghosh

**Library Preparation and Sequencing:** Murshed Hasan Sarkar, Md. Saddam Hossain, Iffat Jahan

**Data Analysis and Primary Analysis:** Sanjana Fatema Chowdhury, Murshed Hasan Sarkar, Syed Muktadir Al Sium, Tanay Chakrovarty

**Manuscript Preparation:** Sanjana Fatema Chowdhury, Murshed Hasan Sarkar, Syed Muktadir Al Sium, Showti Raheel Naser

**Logistic Support:** Md. Ahashan Habib, Shahina Akter, Tanjina Akhtar Banu, Barna Goswami

## Acknowledgements

The authors would like to acknowledge National Institute of Laboratory Medicine & Referral Center (NILMRC), Dhaka, Bangladesh for the support during sample collection. The authors also acknowledge Bangladesh Council of Scientific and Industrial Research (BCSIR), Bangladesh and Ministry of Science and Technology, Bangladesh for the funding and infrastructure support.

